# AYUKA: A toolkit for fast viral genotyping using whole genome sequencing

**DOI:** 10.1101/2022.09.07.506755

**Authors:** José Afonso Guerra-Assunção, Richard Goldstein, Judith Breuer

## Abstract

Technological advances enabled the frequent use of whole genome sequencing in the clinical microbiology laboratory. While generating data is now easier than ever, the computational resources and expertise required for analysis are still a challenge for clinical applications. Since it is not always possible to collect clinical specimens at the peak viral load, sequencing results are also not always amenable for analysis with bioinformatics pipelines that always require high quality data.

Here we present a fast and reliable method, we named AYUKA, for analysis of viral sequencing data that does not require data pre-processing and provides quality control metrics including estimates for sequencing depth and genome coverage, as well as identifying the viral genotypes in a sample and distinguishing mixed infection from recombinants.

This method can be applied to any virus where a classification by genotype is employed and determining it is relevant. We generated a validation dataset composed of cultured and sequenced reference adenoviruses from distinct species, that we compared with the gold standard clinical processing pipeline currently implemented to demonstrate reliability. The validation shows better sensitivity than mapping and perfect specificity in detecting the correct genotypes and in a wide range of adenovirus species. Run time was consistently under one minute per sample on a standard laptop, allowing the analysis of more than 100 samples per hour.

This open-source method is available at https://github.com/afonsoguerra/AYUKA and precomputed databases are available at https://zenodo.org/record/6521576 allowing analysis of raw data straight from the sequencer within minutes on a standard computer, with minimum setup or expertise required to perform the analysis.

The information contained within the AYUKA report can be of use for both the clinical team that collected the sample, but also for guiding the bioinformatics analysis team in the in-depth downstream analyses and genetic epidemiology investigations.

## 2 Introduction

Advances in clinical microbiology have brought to the forefront molecular viral typing and high-sensitivity polymerase chain reaction (PCR) based assays, which form the backbone of viral monitoring protocols. These PCR tests are now commonly followed by high throughput sequencing to acquire more information from the samples of interest, enabling detailed genetic epidemiology studies (Treangen and Pop 2018; Roy et al. 2019; Brown et al. 2019).

There are always a small number of questions that need to be answered before any in-depth analysis. These can relate to the sequencing quality, read depth and genome coverage, but also to the choice of reference genome. This is not always straightforward as there can be multiple genotypes co-existing in the same sample.

The turn-around time for bioinformatics analysis of clinical samples is also important, particularly during a suspected or confirmed viral outbreak. Prompt processing allows rapid monitoring of transmission events and implementation of control measures, as well as making of appropriate clinical decisions.

The surveillance of human adenoviruses, in particular, can benefit from clinical whole genome sequencing. Human adenoviruses are highly diverse, comprising seven species which are subdivided into more than 100 genotypes. Each of the species have slightly different properties and can use different receptors to gain cell access (Lasswitz et al. 2018). Adenovirus infections are quite common in the winter months and can be a cause of the common cold, which is not serious in immunocompetent individuals. Nevertheless, an adenovirus infection can be life threatening for immunocompromised patients who have received an haematopoietic stem cell transplant. This poses great challenges in locations that care for a number of these patients, requiring enhanced cleaning protocols (Pankhurst et al. 2014; Cloutman-Green et al. 2015) and active case monitoring with sequencing of found cases. More recently, human adenovirus genotype F41 has also been implicated into an unexpected outbreak of non-A-E hepatitis in immunocompetent children under the age of 16 in the United Kingdom and in several countries (UKHSA 2022; Morfopoulou et al. 2022).

Genetic epidemiology based on whole genome sequencing has been shown to be an effective approach to identifying transmission links between adenovirus samples (Houldcroft et al. 2018; Myers et al. 2021). Identifying the specific genotype(s) present in a sample can immediately separate the samples that are of a genotype of interest from those that are unrelated to an outbreak under investigation. Conversely, samples that share rare recombination patterns often are part of the same transmission chain. Samples containing mixed infections of different genotypes tend to have a worse prognosis and for these reasons, providing a quick assessment is beneficial for the clinical team.

New adenovirus genotypes commonly arise by lengthy periods of intra-host evolution and mutation accumulation, or by extensive recombination between distinct existing genotypes. These recombination events can occur when the same individual harbours a simultaneous infection of multiple genotypes. This makes it particularly relevant to monitor all samples for mixed infections or transmission of viruses that are in themselves already the product of recombination of different known genotypes.

While traditional adenovirus genotyping focuses on three genes alone and their respective PCR products, the use of WGS brings more detail and allows the inference of transmission events between samples. The existence of a well-supported, fast, streamlined, and easy to use bioinformatics pipeline for viral analysis brings these abilities closer to the bedside where they can have an impact on patient outcomes and in reducing the size of eventual outbreaks, by fast detection of potential outbreaks during routine genotype monitoring. This also allows the prospective analysis of samples is almost real-time.

In this manuscript we introduce AYUKA, a new sequencing analysis method for viral genome analysis. In a few seconds per sample, it performs genotyping, mixed infection, and recombination detection, using raw data produced by a genome sequencer. It was designed with ease of use in mind such that no specialist knowledge would be required to derive clinically actionable insights from WGS data.

The aim of the AYUKA pipeline is not to be an endpoint in itself but rather an advanced but easy-to-use screening method that can simultaneously infer sequence quality (coverage and sequencing depth), viral genotype, and in the cases where multiple viral genotypes are confidently detected, distinguish between multiple infection and recombination. This wealth of data and fast speed makes this an ideal starting point to guide a more detailed reference-based or de novo assembly-based sequence analysis.

AYUKA is particularly relevant during outbreaks, that for adenovirus can commonly occur in winter months, as the genotyping alone can help separate samples that might be part of the outbreak from samples that are from a different genotype.

To achieve speed and accuracy, AYUKA is based on k-mer counting analysis. k-mers are short, fixed length nucleotide sequences, which have been used successfully for a wide variety of biological sequence analysis tasks such as de novo assembly, genome quality control and metagenomics. AYUKA is based around the jellyfish2 (Marçais and Kingsford 2011) k-mer counting library, which is one of the many highly optimised, open-source k-mer libraries popularly used for similar applications. K-mer methods are inherently fast, as they can be implemented using highly optimised string-matching procedures and simple robust statistical tests. We can further speed computations by restricting our analysis to those k-mers that provide the most power to distinguish between virus genotypes.

While common metagenomics software could potentially be configured to perform the genotyping step with some success, these approaches are not be able to distinguish recombination and mixed infections. Because of the much larger database required to run metagenomic searches, the computational resources required are also significantly higher than for AYUKA.

By providing an open-source tool that users can modify to their own needs as well as a fully containerized solution with self-contained dependencies on the software side and databases for different viruses and sequencing parameters on the dataset side, we hope to facilitate the easy adoption in other settings.

## 3 Results

### Algorithm

K-mers are short nucleotide sequences of length k that correspond to substrings of the sequencing data without allowing for mismatches. Because of their fixed length and exact matching, there is a wide variety of efficient algorithms for k-mer counting.

The AYUKA algorithm consists of two distinct steps: database building and sequence classification (see flowchart in Figure 1). The database step consists of the calculation and annotation of k-mers in a relational database (see Materials and Methods). By classifying the k-mers in advance, we can reduce the computation that needs to happen at classification, thus ensuring fast sample classification while maintaining sensitivity and specificity. While the database building step requires more extensive computations, it only needs to be performed if the set of genotypes to analyse or the k-mer length changes, due to sequence updates. We provide a variety of ready-to-use databases for different parameters on the zenodo research hosting platform (https://zenodo.org/record/6521576).

**Figure 1:**
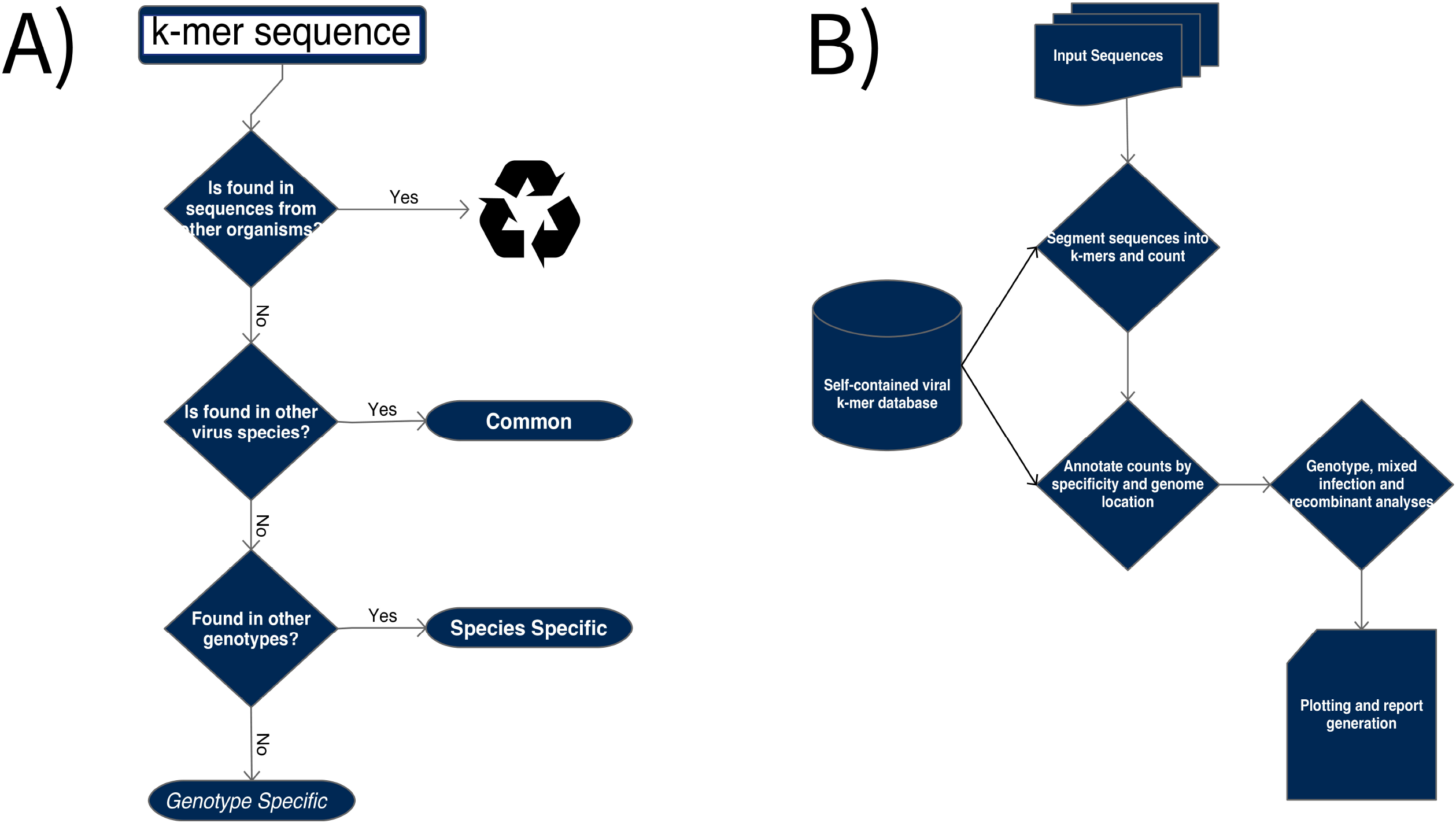
Flow diagram of the AYUKA algorithm. A) Data flow during AYUKA database building steps of the pipeline; B) Data flow for the main algorithm used during AYUKA classification steps, to genotype samples.

The main classification step of the algorithm consists of counting k-mers from the raw reads, cross-referencing with the pre-computed database to classify each detected k-mer, statistical testing to determine which genotype(s) are present, and a step of plotting and report generation (which can be adjusted with the appropriate command line options).

This methodology is sequencing technology independent, provided the sequencing data is in FASTQ or FASTA formats and is also able to analyse consensus sequences if parameters are adjusted to disable the default read depth filters.

The classification step will produce outputs in two main formats, a human readable report in PDF format and a set of tab delimited text files that can be used by subsequent scripts as part as of a mapping or *de novo* assembly-based bioinformatics pipeline.

### Description of the outputs and the report structure

AYUKA produces outputs in two main formats, a human readable report in PDF format and a set of tab delimited text files that can be used by subsequent scripts as part as of a mapping or *de novo* assembly-based bioinformatics pipeline.

The report aims to be a fully versioned, reproducible, and independent result, and contains short explanatory texts to provide guidance on how to interpret the results and how they were computed in this run. For full reproducibility, every report also includes a table indicating the database details, version, and checksum, as well as parameters and thresholds used while running the software. This allows the results to be assessed in context and reproduced later if required.

The first section of the report focuses on the results from the fisher exact test on over-representation of genotype specific k-mers, with a table indicating the genotype(s) found, the GenBank identifier of the representative sequence used for the genotype, the absolute number of different k-mers found in the sample and the corresponding multiple-hypotheses corrected p-value. In case no genotype meets the significance threshold necessary to appear in the table, the report will still be produced but a message will indicate that no statistically significant results were found for the sample and the parameters specified.

If a genotype is confidently found, the second table of the report will show the fraction of genotype specific as well as total k-mers for the genotype that are found in the sample under analysis. This can be a good indicator for how good the coverage is along the genome with that genotype.

Because k-mer counting requires perfect matching between the reference sequence in the database and the sequence data being processed, these values are lower than what is obtained by mapping the data allowing for mismatches and can be useful as a lower-bound estimate for the actual coverage.

An estimate of the read depth along genotype specific k-mers is also provided, which correlates well with the mean read depth before PCR duplication is considered (See Figure 2). Taken together these measures provide a way to assess the data quality available for the sample to proceed for downstream analysis.

**Figure 2:**
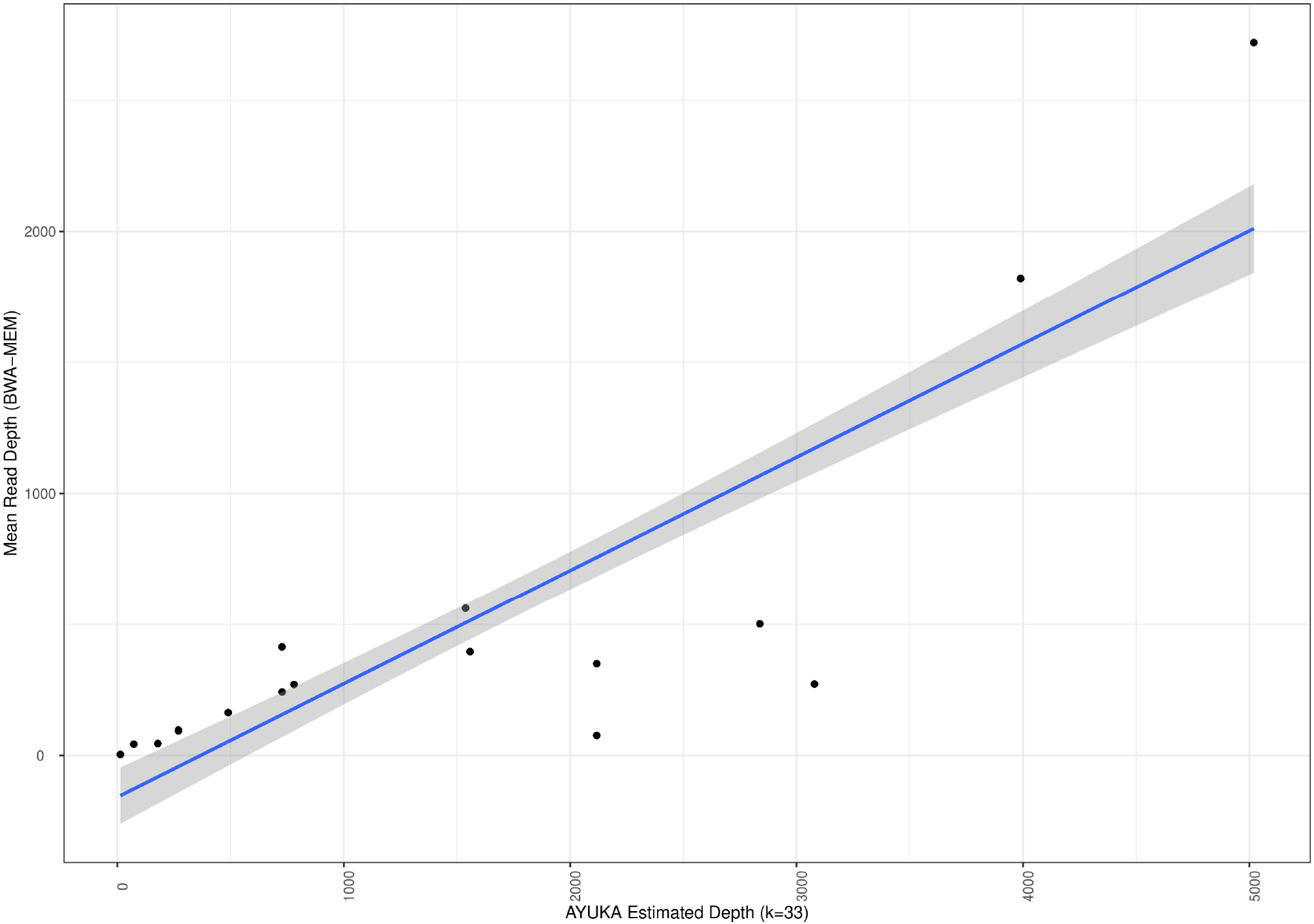
Comparison of AYUKA QC metric estimates with those obtained from reference-based mapping. Scatter plot comparing AYUKA read depth estimates (x axis) with the depth obtained when mapping the data with BWA-MEM to the same genome (y axis). The blue line represents the linear model that best fits all the points and the shaded area its 95% confidence interval. In AYUKA data is used straight from the sequencer and without PCR deduplication, accounting for the higher depth estimates compared with the BWA-MEM values.

### Mixed infection and recombinant definitions

For the cases where more than one genotype are detected, further sections are included to help distinguishing between mixed infections and recombinant viruses in the sample.

Mixed infections are indicated when there are two or more genotypes that present consistent coverage along the whole genome. While there is a possibility of 50/50 mixture of two genotypes, commonly the different genotypes are present in different fractions, which can be seen in the plots and in the estimated depth in the report tables, which should also indicate high coverage values.

Recombination detection relies on genotype-specific k-mers, and assesses if certain areas of the genome have exclusively one of the genotypes. The different genotypes that form part of the recombinant tend to have similar estimated read depths on plots and tables, and low and uneven coverage estimates.

In immunocompromised patients, it is not uncommon to observe the simultaneous presence of a recombinants with a mixed infection. Because of the possibility of complex samples containing both mixed infections and recombinants simultaneously (see Figure 3), and the difficulties of separating these cases statistically, we have chosen to present the data in a graphical way to showcase the proportion of each genotype across the length of the genome, first considering all genotype k-mers (which highlights and allows the inference of the proportions of each genotype in a genomic mixture) and in the second section, we consider only the genotype specific k-mers, which allows for the inference of medium to large recombination events between different genotypes. While recombination events involving short sequences are possible, they are rare and well tolerated by a standard reference-based pipeline downstream of AYUKA.

**Figure 3:**
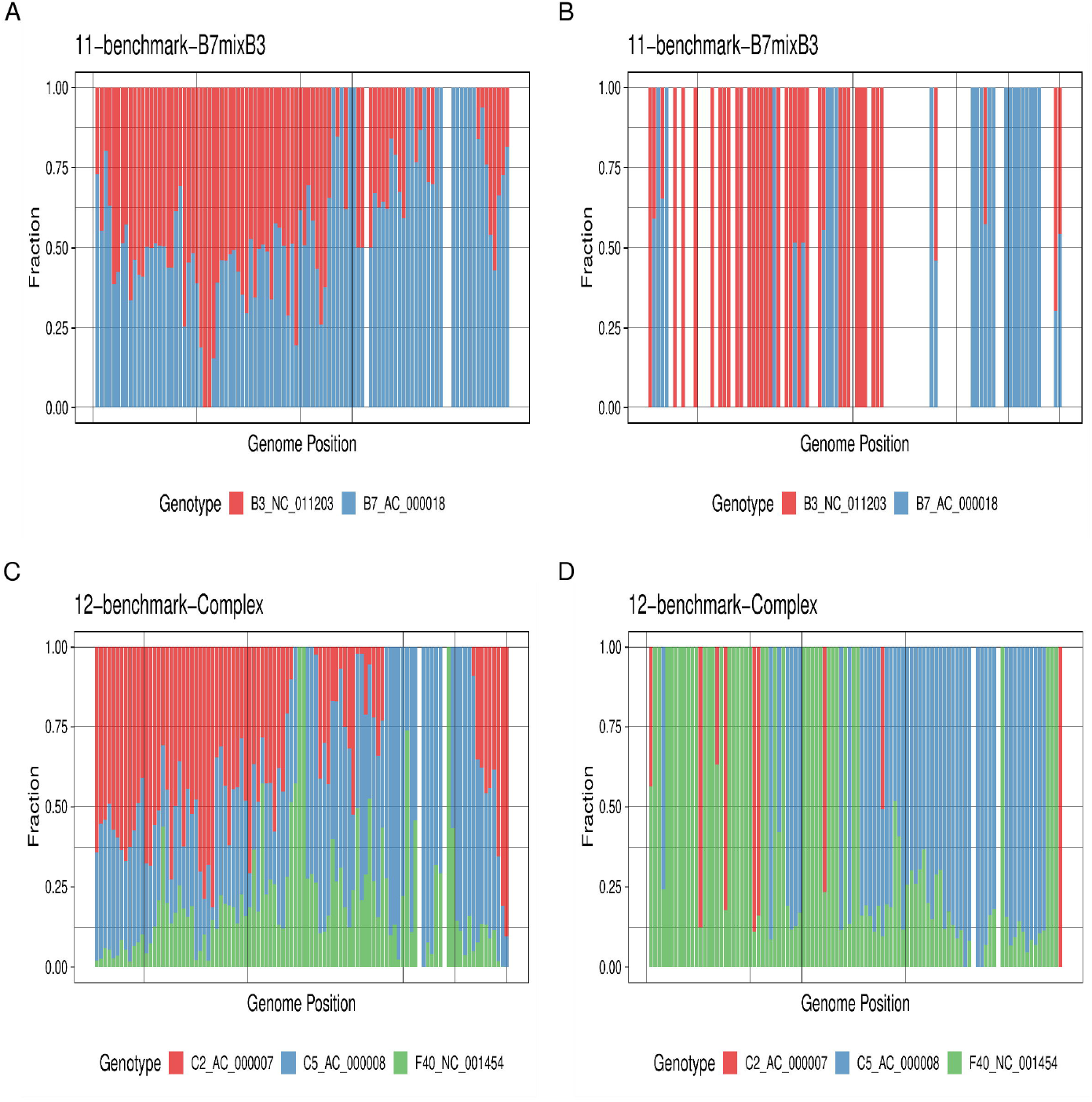
Example of mixture and recombinant identification plots, showing a complex case including a mixture and a recombinant simultaneously. Plots A and B refer to benchmark sample 11, showing fractions of all k-mers (A) or only genotype specific k-mers (B). Areas where there are white gaps represent areas of the genome alignment of all genotypes in the database where there is no coverage for this genotype. Plots C and D show similar information for benchmark sample 12.

An example of how this is visualized is shown on Figure 3, highlighting that some samples encountered in practice, particularly in immunocompromised individuals, can show evidence of multiple events occurring over time and shaping the complexity of the sample observed. This complexity makes it particularly challenging to correctly classify mixture and recombination events statistically, and is better addressed using a specialist pipeline downstream, for example BBSplit (‘BBMap’ 2022) for mixtures and RDP4 (Martin et al. 2015) for recombination analysis when these events are found by AYUKA.

### Control Datasets

To assess the performance of the AYUKA algorithm under different conditions, a set of reference material with known genotype was procured from either ATCC or a local sample biobank, cultured and sequenced (see Materials and Methods and Supplementary Table 1). This dataset also contains one sample that is known to be a mixture and one that has been shown to reflect a mixture of a pure sample and a recombinant. To test method sensitivity, the sample for genotype F41 was down-sampled, using picard (‘Picard Toolkit’ 2019) to obtain fastq files with a lower number of reads.

These reference samples were then processed with AYUKA followed by a full reference-based mapping, PCR duplicate removal, variant calling and consensus building bioinformatics pipeline. The obtained results were then compared and for the basis of this manuscript. The same set was also used for the metagenomics comparison.

This dataset is as diverse as possible, containing at least one genotype from each of the main human adenovirus species (see Supplementary Table 1).

### Sensitivity and Specificity in the context of k-mer length

The sensitivity and specificity of the AYUKA algorithm can be adjusted primarily by the choice of k-mer length.

To test the various effects of k-mer length on the algorithm, we generated databases for adenovirus genotype classification containing references from GenBank (see Supplementary Table 2) in k-mer lengths 17, 33, 39, 65, 89, 139, 239 and 289. These values were chosen to be odd numbers around 10 bp shorter than the most common Illumina read lengths.

While there is no limit in the algorithm for how short or long the k-mer should be, a short k-mer will be non-specific and have little classification value, while k-mers longer than the read-length will not match the reads being analysed.

Because the k-mers are matched exactly to the reads, divergence from the reference sequences in the database will cause the number of detected k-mers to drop, so we see lower sensitivity with higher k values, which can be inferred from the lower depth and coverage estimates obtained (Figure 4).

**Figure 4:**
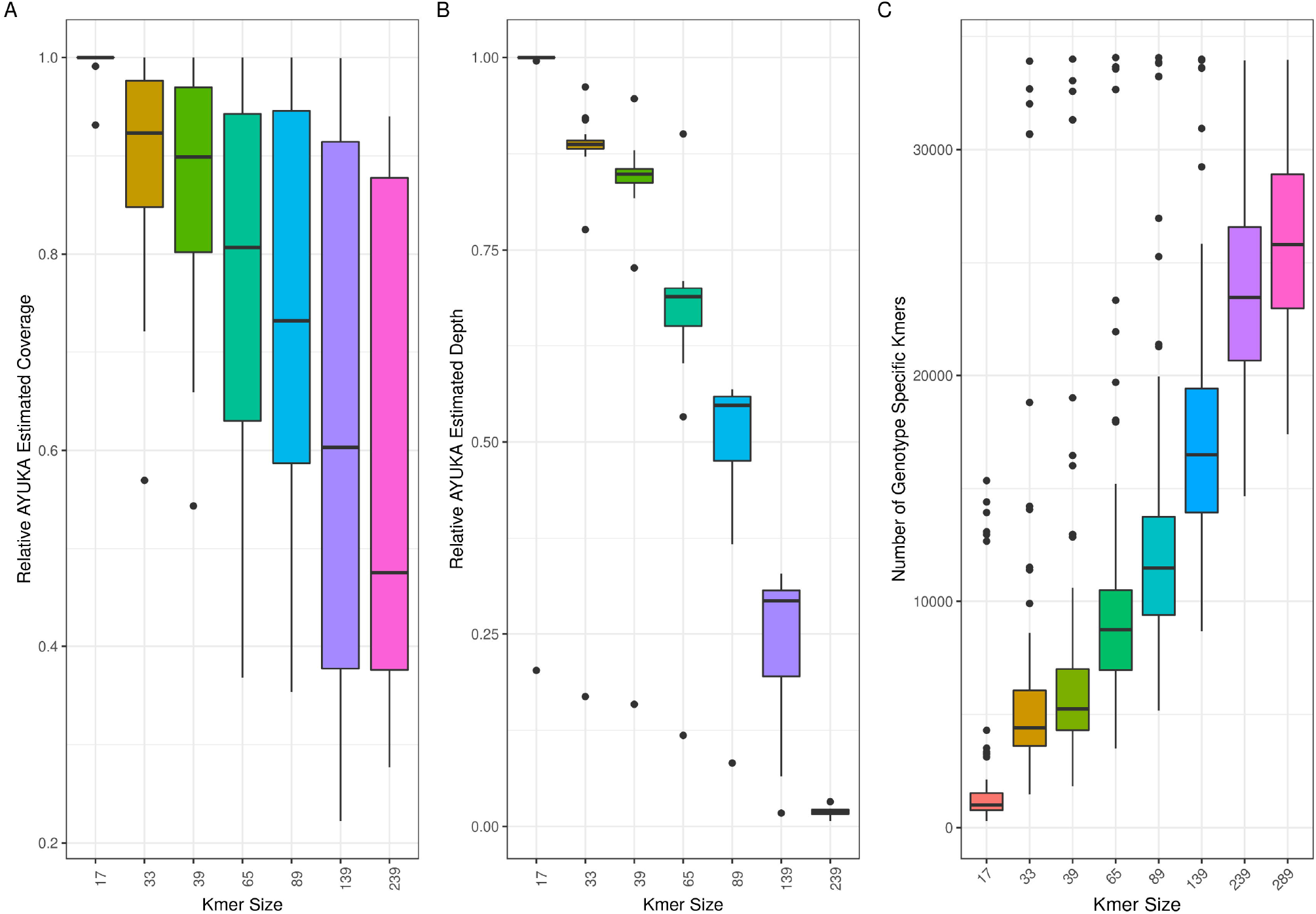
Relationship between choice of k-mer size and AYUKA inferences. For each benchmark sample, the highest estimate for coverage produced by AYUKA for each sample was used as a reference value. This was then compared with the estimate produced at different k-mer lengths for Coverage A) and read depth B), which can be used as a proxy for method sensitivity with varying k-mer lengths. Sensitivity can be estimated by the number of genotype specific k-mers for each genotype present in the database, which is shown in C).

Conversely, longer k-mers are more specific than short k-mers, as they encapsulate more of the genotype specific information in a single sequence. Predictably, we see the number of genotype specific k-mers in the database rise with higher k values (Figure 4C).

In practical use we tend to use k=33 for most analyses, with an increase to higher k values for more complex samples or when higher specificity is needed. Smaller k values are not specific enough for adenovirus but might work well in other contexts.

The suitability of a certain k-value for a particular viral database can be analysed theoretically, by assessing the fraction of genotype specific k-mer sequences compared with the number of k-mers shared between any pair of reference sequences within the database. This analysis also enables the database designer to ascertain if certain pairs of references are too alike to be part of a classification database (see Supplementary Figure 1).

### Speed benchmarks

A crucial aspect for the design of AYUKA was the ability to produce accurate results quickly, without requiring specialist computer equipment. To assess this goal, we benchmarked the software on a four core/eight thread x86 laptop with 16GB of RAM, running the Linux operating system. We have also managed to run it successfully, albeit significantly more slowly on an ARM based Raspberry Pi 4 single board computer with 2GB RAM.

The benchmark involved running each of the eighteen test samples (see Supplementary Table 1) for each of the eight k-mer values, resulting in 144 runs, which completed in 64 minutes, running one sample at a time.

Detailed resource usage per run is shown in Figure 5A. Here we can see that all runs finished in less than one minute, with an average of just over 30 seconds, and without a direct relationship with k-mer size. Maximum memory usage (Figure 5B) grows with the size of the k-mer database, but even for the larger k values it is still under 1GB RAM, allowing it to run on most modern computers.

**Figure 5:**
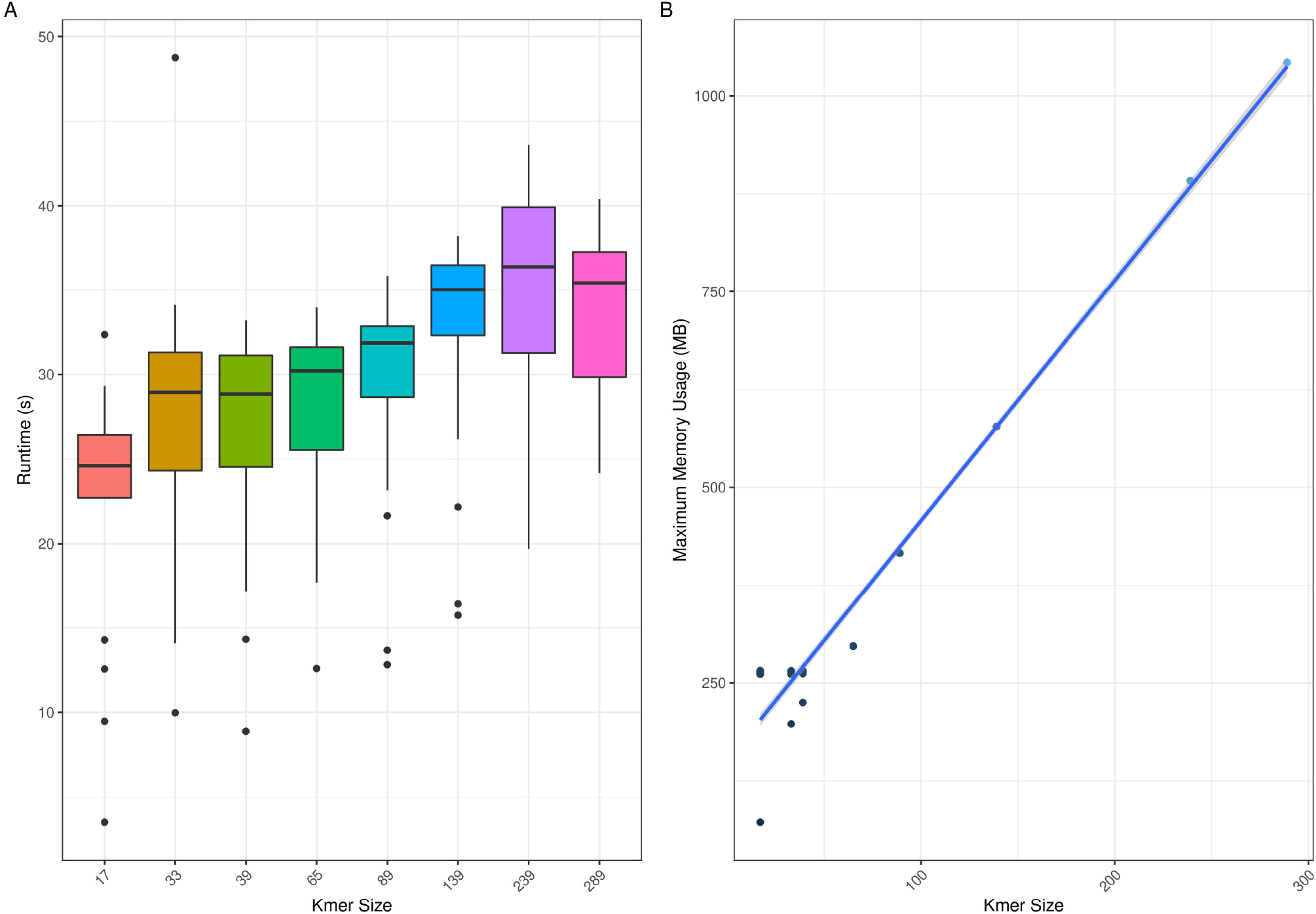
Relationship between k-mer length and computational performance of AYUKA. A) shows the wall-clock run time per sample from the benchmark dataset. B) indicates the maximum RAM usage, in megabytes during the processing of the benchmark samples for each of the k-mer lengths.

A significant amount of time per run is spent generating the figures and report, with performance benefits on running in pipeline mode. This also explains the extremely fast outliers in the plot, corresponding to the few samples where no genotype could be found as such no plots need to be produced.

Like most bioinformatics software, AYUKA relies heavily on input/output operations from the hard drive. Running AYUKA over a network connection or a slow hard drive can have a significant performance impact even if CPU power is available, which impacted our Raspberry pi attempt. Conversely, power-users might want to run it from within a ram-disk.

Speedups can also be obtained by parallelizing, either using multiple CPUs to speed up k-mer counting or by processing multiple samples simultaneously.

### Accuracy benchmarks

The benchmark dataset we put together reflects what is frequently found in real-world samples, in terms of data quality, read depth and diversity. To explore this further we look at three crucial points. We include 10 samples containing a known single genotype, 2 samples containing more than one genotype in more complex situations and 5 samples that were down-sampled in-silico from the F41 genotype sample to assess how the algorithm performs with lower data amounts. We compare these results with results obtained with our standard processing pipeline.

We want to assess the accuracy of the genotype calls, including the ability to handle complex samples, as well as the usefulness and reliability of the estimated QC (Quality Control) metrics provided.

Method sensitivity is superior to other approaches, with 2000 raw reads (which include human, other contaminants, and low-quality reads), sufficient for the samples to be accurately classified as genotype F41 up to k=65 (see Figure 6). The sample with 200 reads did not contain enough depth to be correctly classified with default parameters. In practice, higher sensitivity might be obtained by reducing the read depth requirements at run-time, which would also apply if classification of assembled sequences is desired.

**Figure 6:**
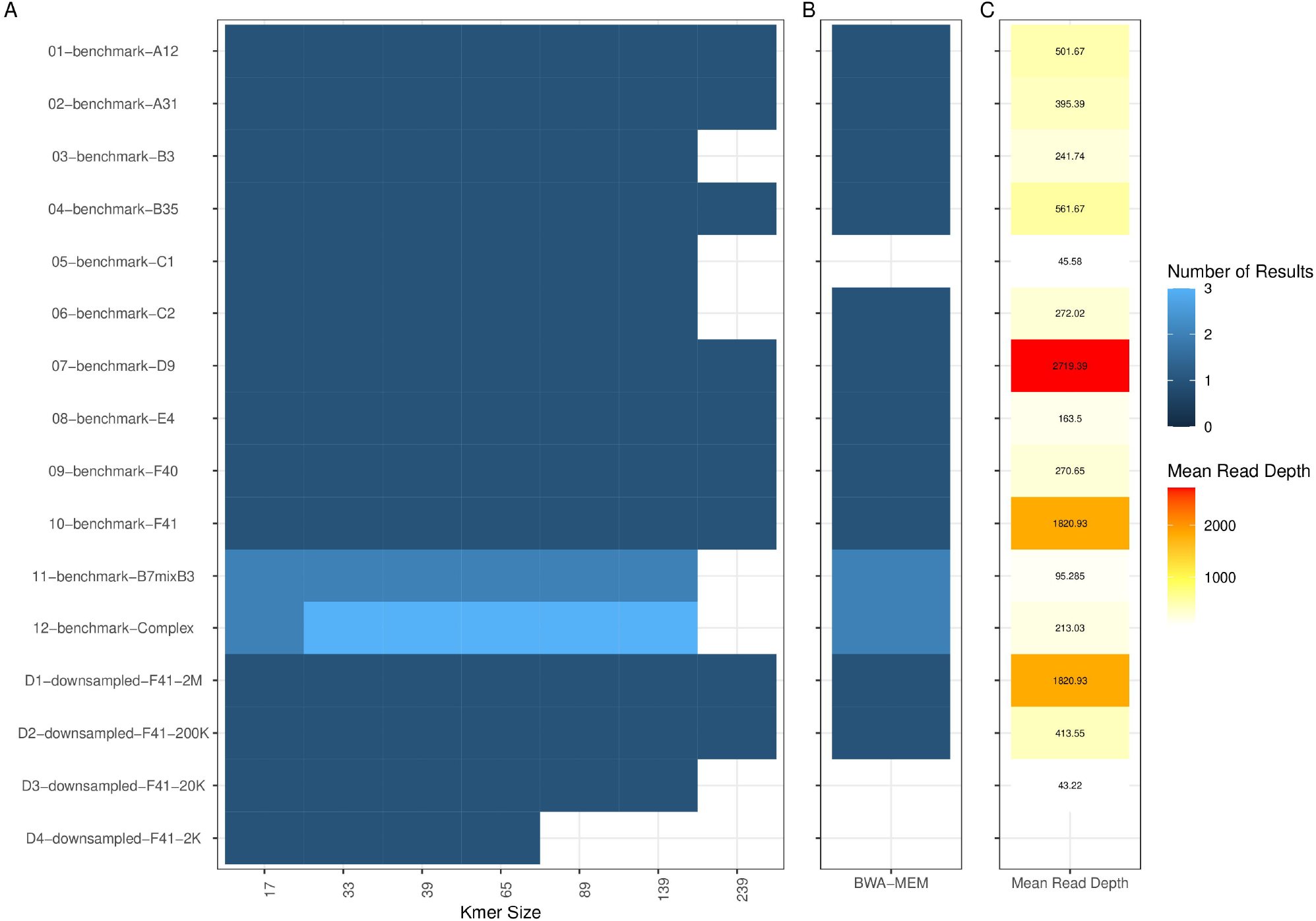
Comparison of results between AYUKA and a reference based viral genome analysis pipeline. A) shows the number of genotypes found to be present in AYUKA results, with results shown for different k-mer lengths. B) For reference-based analysis, we keep samples for phylogenetic analysis if a consensus genome can be obtained with at least 90% coverage at 30X read depth. C) shows the Mean Read Depth obtained in the reference-based analysis for each sample in the dataset. This highlights the wide range of read depths and data quality used to benchmark AYUKA in this context, and its superior sensitivity when compared with a standard reference-based analysis.

We also observe that the correct genotypes are assigned for all other test samples, demonstrating the good specificity that this approach provides across a wide range of mean read depths (Figure 6). By focusing on k-mers with genotype discrimination power, AYUKA achieves a higher specificity than what is obtained with reference mapping methods, but it is also important to note that reference-based consensus building, provided there is enough signal in the data, can correct a closely related reference sequence to a consensus sequence of the sample, as for the most cases there is not an exact matching reference in the databases. Nevertheless, some larger recombination events might still be missed by reference-based methodology and cause areas of spurious variation in genome analysis, disrupting phylogenetic approaches.

We observe a good correlation between the AYUKA estimated sequencing depth and the mapping inferred mean read depth (Figure 2). The value tends to be higher in AYUKA as the data is used raw, without pre-processing while the mapping data has had PCR duplicates removed as well as read quality filtering applied, thus reducing the numbers. The AYUKA estimate still provides a good proxy to distinguish heavily sequenced samples from low read depth samples.

Because of the tendency of k-mers to not be detected when there are sequence divergences from the reference sequence, we find that the coverage estimates are useful within the AYUKA algorithm for the user to assess how much of the database in reflected in the sample, but they do not form a good proxy for what coverage is obtained in mapping, especially at higher k values.

### Comparison with metagenomics

To a certain extent, viral genotyping can be considered a specific case of viral metagenomics. To provide a comparison, we have processed the benchmark samples we generated through the popular kraken2 (Wood, Lu, and Langmead 2019) metagenomics pipeline. To produce a similar output, these results were further processed with the bracken (Lu et al. 2017) companion program. These tools were chosen due to their popularity and use of k-mers as a key part of the algorithm.

The results are shown in Supplementary Table 3. While kraken2 seems to provide some guidance over what the most prevalent genotypes in the sample are, it frequently reports other adenovirus genotypes that were not expected to be present in the sample, as well as other organisms, including the human cells used to grow the viruses. This makes interpretation of the results difficult and less suited for clinical use. The kraken2 results also lack the other components of AYUKA that make it possible to distinguish mixed infections from recombinant samples. The computational resources required to run kraken2 are much higher than for AYUKA, limiting its usability in places that do not have dedicated bioinformatics infrastructure.

## 4 Materials and methods

### Database building

The database building step starts with two text files produced by the user to configure the characteristics of the database being built. The first is a tab delimited two column list of genomes to be included in the database, with the first column detailing the ID for the genotype, and the second column the GenBank identifier for the genotype desired (see Supplementary Table 2). The genotypes chosen should be as distinct as possible and complete, without many ambiguous characters. The second file contains the remaining parameters for the database. Example files and parameters are provided with the code. Once the configuration files are available, the process can be started by using the provided Makefile.

In the first step, the required sequences are retrieved from GenBank, named, and formatted appropriately. This is followed by k-mer counting, using the Jellyfish algorithm, for each file independently as well as the combined set of all sequences, thus defining our k-mer set.

Once all k-mers of interest are identified, the NCBI provided NT database is either streamed directly from the site using cURL or read from a local file (if desired), to identify k-mers that are non-specific to the species of interest. It is necessary to ignore in the NT database all instances of the species of interest. This can be done by a combination of an exclusion string that can be defined in the configuration file or with a list of sequence identifiers to ignore during filtering (e.g. - synthetic constructs that are known to be derived from the virus of interest). After this step, only k-mers that are specific to the virus of interest remain. Because the k-mer counting is restricted to the k-mers originally identified, the memory requirements are reduced, and limited for each database size, with the time being spent mostly on I/O operations (see Figure 5).

With the final k-mer set defined, we then pre-compute all values that can be useful for the later statistics, including the total number of k-mers per genotype, the number of genotype specific k-mers per genotype, and the position of each k-mer in each genome of interest. The positions are also binned in bins corresponding to 1% of the genome, for noise reduction and ease of plotting.

The next step, that can be disabled if not of interest, consists of generating a multiple sequence alignment using the G-INS-I algorithm available in the MAFFT package, encoding of the alignment as a relational database with the coordinates of all the genomes of interest and conversion of the k-mer positions and bin-boundaries to alignment coordinates, which is followed by pre-computing of the positional statistics for each bin.

The last step consists of saving all the different tables, as well as database metadata and timestamp in a SQlite3 relational database that can then be used by AYUKA for sample analysis.

### Classification algorithm

The AYUKA Software is implemented in the PERL programming language, with the use of R for plotting and LaTeX for report generation.

The classification algorithm starts by loading the database of interest and identifying the total k-mer set that is contained within. The user can classify sequences that are stored in either FASTA or FASTQ file formats. The data within can be raw read data, contigs from de novo assembly analysis or consensus sequence from a reference-based analysis. The provided sequences are read using Jellyfish to produce counts for each of the k-mers in the database. A read depth threshold is employed for performance, and to reduce sequencing noise. By default, this threshold has a value of 10. If the sample is of particularly low depth, or the input data is not raw data but an already processed sequence, this threshold can be removed as a command line parameter (--min_depth).

Once the k-mers are counted, the data is linked to the annotations in the database to assess how many k-mers, which class and position they belong to.

Some viruses are very prone to recombination and result in small fragments of different genotypes along the genome. Since the objective is to find the major genotypes that are present in a sample, there is also a threshold that can be used to set a minimum of the genome that should be covered by a certain genotype before it is considered a valid result. By default, this is 0.05 and can be set with the parameter (--min_geno_frac).

A Fisher exact test, with corrections for k-mer independence and Bonferroni multiple test correction is then employed to determine which genotypes are present in the sample above the expected sequencing noise level. It is important to note that the AYUKA sensitivity is higher than that of the common downstream pipelines, so a positive genotype identification does not guarantee success in the downstream analyses.

If no genotype is found to pass the statistical test, the analysis will conclude with the production of a report that will state that there were no significant results and the parameters used during the analysis, for reproducibility.

If there is at least one candidate genotype for the sample, the count tables are processed using the R Statistical Programming language to produce plots and tables for the report.

Following statistical testing, the first results focus on coverage and depth estimates. If more than one genotype is present, a plot will be generated for the fraction of total k-mers (containing both k-mers that are specific to the genotype as well as k-mers that are shared between genotypes) as they are present along the genome. This should allow easy identification of mixture fractions along the genome if that is the case. This is followed by a similar plot containing just genotype specific k-mers to aid visual inspection of recombinant samples.

Once all data for the tables and plots is computed, the PDF report is produced using pdflatex from the LaTeX distribution.

The algorithm is flexible enough that the user can choose to start with an already existing jellyfish file of counted k-mers (but is responsible for checking that it matches the database being provided) and skip report production or even statistical testing. These modes are useful if AYUKA is fully integrated as part of an end-to-end processing pipeline that contains other checks and produces its own reports, and only parts of the AYUKA algorithm are deemed required.

### Running the software

The software package has several dependencies. As such, the recommended way to execute the algorithm is through the singularity container available in the GitHub releases at https://github.com/afonsoguerra/AYUKA or that can be generated by the provided Singularity recipe file. This file can also be used as guidance for the installation of all required dependencies in an Ubuntu Linux based system.

In testing we found the use of containers does not affect algorithm performance and is of great advantage for deployment in the cloud or in HPC (High Performance Computing) systems where super-user access is not available.

### Benchmarks

A selection comprising at least one example of adenovirus species A to F was procured from the ATCC standards from existing control samples from and cultured. From the cultured samples we produced libraries using the Agilent SureSelect protocol as previously described (Houldcroft et al. 2018). This reproduces the protocol developed in our lab for processing clinical samples. See Supplementary Table 1 for details of the samples used in this positive control dataset.

To ensure that the algorithm is impervious to noise, we produced dinucleotide shuffled versions of sequences for each genotype in the database using uShuffle (Jiang et al. 2008) and processed them using AYUKA, to ensure no false positives were likely to occur when using the default parameters and the Fisher exact test is appropriate for controlling noisy data.

To compare the data inferred by AYUKA, the positive control samples were processed using the standard pipeline. Following a first step of quality control and adapter trimming, the data is genotyped by comparing the best mapping from the first mapping step against all references simultaneously with high specificity parameters. After genotyping, the trimmed read data is mapped using bwa mem to the relevant reference sequence.

To compare the genotypes inferred by AYUKA with a standard metagenomics tool that is also k-mer based, we analysed out positive control samples using kraken2 (Wood, Lu, and Langmead 2019) and aggregated the data into species calls using the companion software bracken (Lu et al. 2017). The unconstrained standard database was retrieved from https://benlangmead.github.io/aws-indexes/k2 and used with default parameters.

## Discussion

The use of whole genome sequencing for the analysis of viral outbreaks is a promising approach that has shown superior results for infection prevention and control. The widespread application of such techniques is still hampered by the lack of resources for the analysis of such datasets. Here we present a method that has low computational requirements, does not require advanced expertise to operate, and improves the efficiency of existing pipelines by determining within a few seconds the genotype composition of a sample containing the virus of interest.

This method acts as an initial screening step for a more complex pipeline, saving time and resources in deciding what genome reference to use, and helping to exclude low quality samples or flag complex samples for more in depth analysis. The extensive annotation of the versions and databases used for analysis enables the perfect reproducibility of any results obtained. Combined with versioned hosting at zenodo, this should provide good reproducibility in the long term.

Most of the advantages of the AYUKA methodology were highlighted in the investigation we undertook to explore the causes of the unexpected outbreak of acute hepatitis of unknown aetiology in children in early 2022 (Morfopoulou et al. 2022; UKHSA 2022). The high sensitivity enabled us to detect virus and reliably genotype several samples where the virus had been picked up by PCR and/or metagenomics, but not enough depth was available for a more thorough variant analysis.

After the successful validation of the method using adenovirus samples of known genotype, we present in this manuscript, we expanded the database set producing databases for human adeno-associated viruses (AAV), as well as Epstein-Barr virus (EBV). The results from these agreed with other research tools for those viruses and support the wide applicability of the AYUKA method in other contexts not presented here.

The fast speed of the pipeline and ease of use make it well suited for contamination control in RNA-seq analyses, or for the development of custom databases to control payload insertion in genetic engineering experiments involving Adenovirus, AAV or other viruses as vectors.

Even though AYUKA shares one of the limitations of metagenomics, in which only genotypes that are pre-selected to integrate the database are capable of being detected, this method is not meant as a replacement for metagenomics approaches, as it requires the user to have some prior knowledge of the virus they would want to genotype.

We also do not envision AYUKA being used as a terminal endpoint in an analysis pipeline, as more insights can be obtained from the data by performing an in-depth reference-based or *de novo* assembly-based bioinformatics analyses, if resources, expertise and data quality permit such an analysis.

In this approach, sensitivity and specificity are mostly affected by the choice of k-mer size. Larger k-mers have better ability to distinguish genotypes that are genetically quite similar, but because this method relies on exact matching to the reference sequence database, large k-mers have reduced sensitivity, especially for samples that are very divergent from the reference database. It is also important to note that the k-mer length must always be smaller than the read length of the raw data being analysed.

This approach was developed to solve a specific problem, but we have found it invaluable in other contexts and for other viruses. Low resource requirements mean it can be used by anyone even if they do not have a massive computer. With the capacity to analyse more than one hundred samples an hour, it is suitable for integration in many pipelines without a large performance penalty. We hope that providing it as a containerized solution will ease its adoption and promote its use. We look forward to expanding the range of databases available for other viruses.

## Acknowledgements

J.A.G.-A. Wishes to thank the GOSH (Great Ormond Street Hospital) infection prevention and control team for constructive feedback, Dr. Cristina Venturini for testing out the algorithm with EBV, and the Breuer Lab and UKHSA for constructive feedback. The authors acknowledge the use of the UCL Myriad High Performance Computing Facility (Myriad@UCL), and associated support services, in the completion of this work.

