## Supplementary Material for "AYUKA: A toolkit for fast viral genotyping using whole genome sequencing"

### Supplementary Information

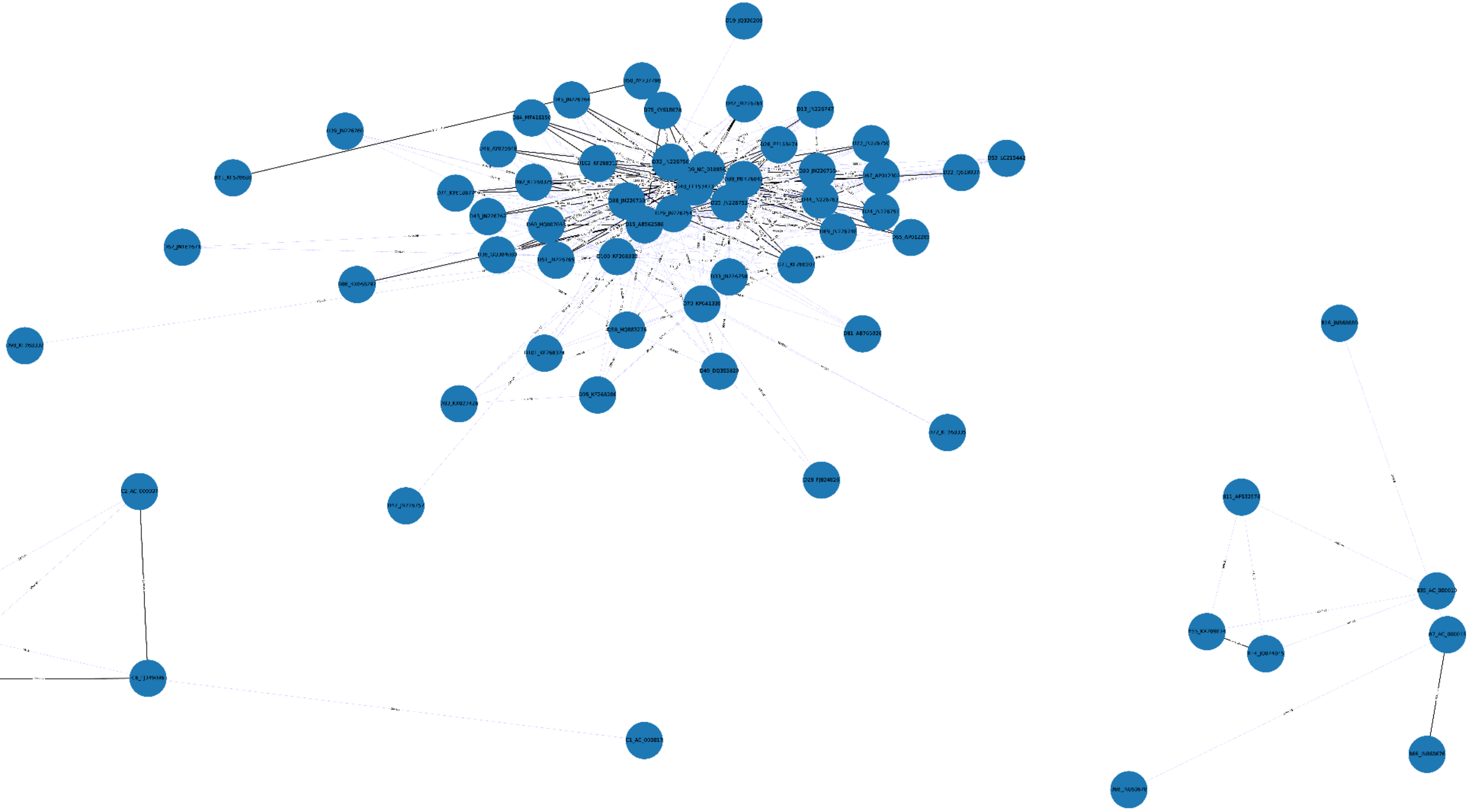

**Supplementary Figure 1:** Graph linking different genotypes by shared k-mer sets. Pairwise k-mer distances were computed from the AYUKA database with k=33. Genotypes that are fairly distinct from others are not shown. Tenuous k-mer overlap is shown in light blue lines, while strong similarity is shown by a black line. Links between distinct genotypes are not shown.

**Supplementary Table 1:** Samples in Benchmark Dataset

| Sample Name | Sample Type | Genotype(s) | NumTotalReads |
|---|---|---|---|
| 01-benchmark-A12 | Pure-Genotype | A12 | |
| 02-benchmark-A31 | Pure-Genotype | A31 | |
| 03-benchmark-B3 | Pure-Genotype | B3 | |
| 04-benchmark-B35 | Pure-Genotype | B35 | |
| 05-benchmark-C1 | Pure-Genotype | C1 | |
| 06-benchmark-C2 | Pure-Genotype | C2 | |
| 07-benchmark-D9 | Pure-Genotype | D9 | |
| 08-benchmark-E4 | Pure-Genotype | E4 | |
| 09-benchmark-F40 | Pure-Genotype | F40 | |
| 10-benchmark-F41 | Pure-Genotype | F41 | |
| 11-benchmark-B7mixB3 | Multi-Genotype | B7,B3 | |
| 12-benchmark-Complex | Multi-Genotype | F40,C2,C5 | |
| D1-downsampled-F41-2M | Downsampled F41 Genotype | F41 | 2000000 |
| D2-downsampled-F41-200K | Downsampled F41 Genotype | F41 | 200000 |
| D3-downsampled-F41-20K | Downsampled F41 Genotype | F41 | 20000 |
| D4-downsampled-F41-2K | Downsampled F41 Genotype | F41 | 2000 |
| D5-downsampled-F41-200 | Downsampled F41 Genotype | F41 | 200 |

**Supplementary Table 2:** Samples in AYUKA database

| Adenovirus Species | Genotype | GenBank Accession |
|---|---|---|
| A | A12 | NC_001460 |
| A | A18 | GU191019 |
| A | A31 | AM749299 |
| A | A61 | JF964962 |
| B | B11 | AF532578 |
| B | B14 | JQ824845 |
| B | B16 | JN860680 |
| B | B21 | KF528688 |
| B | B35 | AC_000019 |
| B | B3 | NC_011203 |
| B | B50 | AY737798 |
| B | B55 | KX289874 |
| B | B66 | JN860676 |
| B | B68 | JN860678 |
| B | B7 | AC_000018 |
| C | C1 | AC_000017 |
| C | C2 | AC_000007 |
| C | C57 | HQ003817 |
| C | C5 | AC_000008 |
| C | C6 | FJ349096 |
| C | C89 | MH121097 |
| D | D100 | KF268330 |
| D | D101 | KF268324 |
| D | D102 | KF268312 |
| D | D13 | JN226747 |
| D | D15 | AB562586 |
| D | D17 | AC_000006 |
| D | D19 | JQ326209 |
| D | D20 | JN226749 |
| D | D22 | FJ619037 |
| D | D23 | JN226750 |
| D | D24 | JN226751 |
| D | D25 | JN226752 |
| D | D26 | EF153474 |

| | | |
|---|---|---|
| D | D28 | FJ824826 |
| D | D29 | JN226754 |
| D | D30 | JN226755 |
| D | D32 | JN226756 |
| D | D33 | JN226758 |
| D | D36 | GQ384080 |
| D | D38 | JN226759 |
| D | D39 | JN226760 |
| D | D42 | JN226761 |
| D | D43 | JN226762 |
| D | D44 | JN226763 |
| D | D45 | JN226764 |
| D | D46 | AY875648 |
| D | D47 | JN226757 |
| D | D48 | EF153473 |
| D | D49 | DQ393829 |
| D | D51 | JN226765 |
| D | D53 | LC215442 |
| D | D54 | NC_012959 |
| D | D58 | HQ883276 |
| D | D60 | HQ007053 |
| D | D62 | JN162671 |
| D | D64 | JQ326206 |
| D | D65 | AP012285 |
| D | D67 | AP012302 |
| D | D69 | JN226748 |
| D | D70 | KP641339 |
| D | D71 | KF268207 |
| D | D72 | KF268335 |
| D | D74 | KY618677 |
| D | D75 | KY618678 |
| D | D81 | AB765926 |
| D | D83 | KX827426 |
| D | D84 | MF416150 |
| D | D86 | KX868297 |
| D | D88 | MF476842 |

| | | |
|---|---|---|
| D | D8 | AB746853 |
| D | D92 | KF268325 |
| D | D95 | KF268206 |
| D | D98 | KF268332 |
| D | D9 | NC_010956 |
| E | E4 | NC_003266 |
| F | F40 | NC_001454 |
| F | F41 | MG925782 |
| G | G52 | DQ923122 |

**Supplementary Table3:** Kraken2/Bracken Results

| SampleName | Species Name | Taxonomy ID | Num Reads | Fraction Total |
|---|---|---|---|---|
| 01-benchmark-A12 | Xanthomonas euvesicatoria pv. alfalfae | 359387 | 67 | 0.11424 |
| 01-benchmark-A12 | Streptomyces lividans 1326 | 1200984 | 100 | 0.17003 |
| 01-benchmark-A12 | Human adenovirus 54 | 651580 | 92 | 0.15567 |
| 01-benchmark-A12 | Proteus phage VB_PmiS-Isfahan | 1969841 | 331 | 0.56007 |
| 02-benchmark-A31 | Streptomyces lividans 1326 | 1200984 | 7021 | 0.29609 |
| 02-benchmark-A31 | Xanthomonas euvesicatoria pv. alfalfae | 359387 | 898 | 0.0379 |
| 02-benchmark-A31 | Human adenovirus B2 | 565303 | 1782 | 0.07518 |
| 02-benchmark-A31 | Simian adenovirus 16 | 1715778 | 1524 | 0.06428 |
| 02-benchmark-A31 | Simian adenovirus 3 | 38420 | 5901 | 0.24888 |
| 02-benchmark-A31 | Human adenovirus 5 | 28285 | 655 | 0.02763 |
| 02-benchmark-A31 | Simian mastadenovirus WIV19 | 1923976 | 621 | 0.02619 |
| 02-benchmark-A31 | Bat mastadenovirus WIV12 | 1788434 | 616 | 0.02598 |
| 02-benchmark-A31 | Simian adenovirus DM-2014 | 1560346 | 486 | 0.0205 |
| 02-benchmark-A31 | Simian adenovirus 13 | 38432 | 1602 | 0.06758 |
| 02-benchmark-A31 | Simian adenovirus 25 | 175567 | 356 | 0.01505 |
| 02-benchmark-A31 | Bat mastadenovirus WIV9 | 1788436 | 103 | 0.00434 |
| 02-benchmark-A31 | Simian adenovirus 20 | 585059 | 53 | 0.00228 |
| 02-benchmark-A31 | Human adenovirus 54 | 651580 | 1449 | 0.06113 |
| 02-benchmark-A31 | Staphylococcus phage Andhra | 1958907 | 50 | 0.00211 |
| 02-benchmark-A31 | Proteus phage VB_PmiS-Isfahan | 1969841 | 590 | 0.02488 |

| | | | | |
|---|---|---|---|---|
| 03-benchmark-B3 | Xanthomonas euvesicatoria pv. alfalfae | 359387 | 115 | 0.00133 |
| 03-benchmark-B3 | Streptomyces lividans 1326 | 1200984 | 123 | 0.00143 |
| 03-benchmark-B3 | Human adenovirus B1 | 565302 | 81783 | 0.94237 |
| 03-benchmark-B3 | Human adenovirus B2 | 565303 | 4199 | 0.04839 |
| 03-benchmark-B3 | Proteus phage VB_PmiS-Isfahan | 1969841 | 563 | 0.00649 |
| 04-benchmark-B35 | Xanthomonas euvesicatoria pv. alfalfae | 359387 | 62 | 0.00041 |
| 04-benchmark-B35 | Streptomyces lividans 1326 | 1200984 | 205 | 0.00134 |
| 04-benchmark-B35 | Human adenovirus B2 | 565303 | 152551 | 0.99341 |
| 04-benchmark-B35 | Human adenovirus B1 | 565302 | 240 | 0.00157 |
| 04-benchmark-B35 | Proteus phage VB_PmiS-Isfahan | 1969841 | 503 | 0.00328 |
| 05-benchmark-C1 | Shigella boydii CDC 3083-94 | 344609 | 1575 | 0.0672 |
| 05-benchmark-C1 | Salmonella enterica subsp. enterica | 59201 | 2804 | 0.11962 |
| 05-benchmark-C1 | Xanthomonas euvesicatoria pv. alfalfae | 359387 | 122 | 0.00522 |
| 05-benchmark-C1 | Streptomyces lividans 1326 | 1200984 | 114 | 0.00489 |
| 05-benchmark-C1 | Human adenovirus 1 | 10533 | 16393 | 0.69934 |
| 05-benchmark-C1 | Human adenovirus 5 | 28285 | 2405 | 0.1026 |
| 05-benchmark-C1 | Proteus phage VB_PmiS-Isfahan | 1969841 | 14 | 0.00062 |
| 05-benchmark-C1 | Human bocavirus 2c PK | 1511882 | 12 | 0.00051 |
| 06-benchmark-C2 | Neisseria lactamica 020-06 | 489653 | 422 | 0.00093 |
| 06-benchmark-C2 | Neisseria meningitidis serogroup C | 135720 | 159 | 0.00035 |
| 06-benchmark-C2 | Neisseria meningitidis alpha14 | 662598 | 43 | 0.0001 |
| 06-benchmark-C2 | Neisseria sp. oral taxon 014 str. F0314 | 641149 | 152 | 0.00034 |
| 06-benchmark-C2 | Neisseria elongata subsp. glycolytica | 88719 | 49 | 0.00011 |
| 06-benchmark-C2 | Haemophilus influenzae 2019 | 1232659 | 448 | 0.00099 |
| 06-benchmark-C2 | Haemophilus parainfluenzae T3T1 | 862965 | 683 | 0.0015 |
| 06-benchmark-C2 | Xanthomonas euvesicatoria pv. alfalfae | 359387 | 15 | 0.00003 |
| 06-benchmark-C2 | Campylobacter concisus 13826 | 360104 | 31 | 0.00007 |
| 06-benchmark-C2 | Rothia mucilaginosa DY-18 | 680646 | 500 | 0.0011 |
| 06-benchmark-C2 | Corynebacterium argentoratense DSM 44202 | 1348662 | 168 | 0.00037 |
| 06-benchmark-C2 | Streptococcus mitis B6 | 365659 | 390 | 0.00086 |
| 06-benchmark-C2 | Streptococcus parasanguinis FW213 | 1114965 | 36 | 0.00008 |
| 06-benchmark-C2 | Streptococcus parasanguinis ATCC 15912 | 760570 | 17 | 0.00004 |
| 06-benchmark-C2 | Streptococcus pseudopneumoniae IS7493 | 1054460 | 492 | 0.00108 |
| 06-benchmark-C2 | Prevotella scopos JCM 17725 | 1236518 | 32 | 0.00007 |
| 06-benchmark-C2 | Alistipes onderdonkii subsp. vulgaris | 2585117 | 132 | 0.00029 |
| 06-benchmark-C2 | Alistipes finegoldii DSM 17242 | 679935 | 16 | 0.00004 |

| | | | | |
|---|---|---|---|---|
| 06-benchmark-C2 | Bacteroides vulgatus ATCC 8482 | 435590 | 113 | 0.00025 |
| 06-benchmark-C2 | Human adenovirus 5 | 28285 | 252216 | 0.55497 |
| 06-benchmark-C2 | Human adenovirus 1 | 10533 | 198345 | 0.43643 |
| 07-benchmark-D9 | Human adenovirus 54 | 651580 | 628427 | 0.99966 |
| 07-benchmark-D9 | Porcine adenovirus 3 | 35265 | 115 | 0.00018 |
| 07-benchmark-D9 | Norovirus GII | 122929 | 38 | 0.00006 |
| 07-benchmark-D9 | Norwalk-like virus | 95340 | 15 | 0.00002 |
| 07-benchmark-D9 | Hepatitis C virus genotype 1 | 41856 | 27 | 0.00004 |
| 07-benchmark-D9 | Hepatitis C virus genotype 3 | 356114 | 18 | 0.00003 |
| 08-benchmark-E4 | Xanthomonas euvesicatoria pv. alfalfae | 359387 | 106 | 0.00201 |
| 08-benchmark-E4 | Streptomyces lividans 1326 | 1200984 | 275 | 0.00518 |
| 08-benchmark-E4 | Simian adenovirus 25 | 175567 | 32208 | 0.6063 |
| 08-benchmark-E4 | Chimpanzee adenovirus Y25 | 1123958 | 20532 | 0.38651 |
| 09-benchmark-F40 | Xanthomonas euvesicatoria pv. alfalfae | 359387 | 181 | 0.25637 |
| 09-benchmark-F40 | Human adenovirus B2 | 565303 | 43 | 0.06091 |
| 09-benchmark-F40 | Proteus phage VB_PmiS-Isfahan | 1969841 | 482 | 0.68272 |
| 10-benchmark-F41 | Xanthomonas euvesicatoria pv. alfalfae | 359387 | 11122 | 0.11044 |
| 10-benchmark-F41 | Streptomyces lividans 1326 | 1200984 | 20779 | 0.20634 |
| 10-benchmark-F41 | Mycobacterium tuberculosis variant caprae | 115862 | 27 | 0.00027 |
| 10-benchmark-F41 | Simian adenovirus 18 | 909210 | 16581 | 0.16464 |
| 10-benchmark-F41 | Simian adenovirus 16 | 1715778 | 22067 | 0.21912 |
| 10-benchmark-F41 | Simian adenovirus 1 | 310540 | 18913 | 0.18781 |
| 10-benchmark-F41 | Human adenovirus 5 | 28285 | 202 | 0.00201 |
| 10-benchmark-F41 | Simian mastadenovirus WIV19 | 1923976 | 2654 | 0.02636 |
| 10-benchmark-F41 | Simian adenovirus 25 | 175567 | 1212 | 0.01204 |
| 10-benchmark-F41 | Human adenovirus B2 | 565303 | 1607 | 0.01596 |
| 10-benchmark-F41 | Simian adenovirus DM-2014 | 1560346 | 1356 | 0.01346 |
| 10-benchmark-F41 | Bat mastadenovirus WIV9 | 1788436 | 879 | 0.00873 |
| 10-benchmark-F41 | Human adenovirus 54 | 651580 | 858 | 0.00852 |
| 10-benchmark-F41 | Tree shrew adenovirus 1 | 47680 | 830 | 0.00824 |
| 10-benchmark-F41 | Simian adenovirus 49 | 995022 | 1332 | 0.01323 |
| 10-benchmark-F41 | Simian adenovirus 20 | 585059 | 133 | 0.00132 |
| 10-benchmark-F41 | Fowl aviadenovirus 5 | 172861 | 59 | 0.00059 |
| 10-benchmark-F41 | Staphylococcus phage Andhra | 1958907 | 93 | 0.00092 |
| 11-benchmark-B7mixB3 | Xanthomonas euvesicatoria pv. alfalfae | 359387 | 45 | 0.00166 |
| 11-benchmark-B7mixB3 | Streptomyces lividans 1326 | 1200984 | 206 | 0.0075 |

| | | | | |
|---|---|---|---|---|
| 11-benchmark-B7mixB3 | Human adenovirus B1 | 565302 | 25211 | 0.91575 |
| 11-benchmark-B7mixB3 | Human adenovirus B2 | 565303 | 1496 | 0.05436 |
| 11-benchmark-B7mixB3 | Proteus phage VB_PmiS-Isfahan | 1969841 | 571 | 0.02074 |
| 12-benchmark-Complex | Parabacteroides distasonis ATCC 8503 | 435591 | 461254 | 0.45514 |
| 12-benchmark-Complex | Phocaeicola salanitronis DSM 18170 | 667015 | 20593 | 0.02032 |
| 12-benchmark-Complex | Bacteroides dorei CL03T12C01 | 997877 | 827 | 0.00082 |
| 12-benchmark-Complex | Bacteroides vulgatus ATCC 8482 | 435590 | 32 | 0.00003 |
| 12-benchmark-Complex | Bacteroides fragilis 638R | 862962 | 11667 | 0.01151 |
| 12-benchmark-Complex | Bacteroides fragilis YCH46 | 295405 | 2524 | 0.00249 |
| 12-benchmark-Complex | Paraprevotella xylaniphila YIT 11841 | 762982 | 1941 | 0.00192 |
| 12-benchmark-Complex | Prevotella intermedia ATCC 25611 = DSM 20706 | 1122984 | 148 | 0.00015 |
| 12-benchmark-Complex | Prevotella scopos JCM 17725 | 1236518 | 575 | 0.00057 |
| 12-benchmark-Complex | Prevotella denticola F0289 | 767031 | 41 | 0.00004 |
| 12-benchmark-Complex | Porphyromonas gingivalis AJW4 | 1403336 | 923 | 0.00091 |
| 12-benchmark-Complex | Porphyromonas asaccharolytica DSM 20707 | 879243 | 777 | 0.00077 |
| 12-benchmark-Complex | Alistipes finegoldii DSM 17242 | 679935 | 279 | 0.00028 |
| 12-benchmark-Complex | Alistipes shahii WAL 8301 | 717959 | 128 | 0.00013 |
| 12-benchmark-Complex | Alistipes onderdonkii subsp. vulgaris | 2585117 | 300 | 0.0003 |
| 12-benchmark-Complex | Barnesiella viscericola DSM 18177 | 880074 | 37 | 0.00004 |
| 12-benchmark-Complex | Escherichia coli O157:H7 | 83334 | 63079 | 0.06224 |
| 12-benchmark-Complex | Escherichia coli UM146 | 869729 | 37469 | 0.03697 |
| 12-benchmark-Complex | Escherichia coli APEC IMT5155 | 1329907 | 13228 | 0.01305 |
| 12-benchmark-Complex | Escherichia coli O157:H16 | 930406 | 13012 | 0.01284 |
| 12-benchmark-Complex | Escherichia coli O44:H18 | 861906 | 2178 | 0.00215 |
| 12-benchmark-Complex | Escherichia coli ED1a | 585397 | 1423 | 0.0014 |
| 12-benchmark-Complex | Escherichia coli Nissle 1917 | 316435 | 11134 | 0.01099 |
| 12-benchmark-Complex | Escherichia coli CFT073 | 199310 | 8222 | 0.00811 |
| 12-benchmark-Complex | Escherichia coli K-12 | 83333 | 26972 | 0.02662 |
| 12-benchmark-Complex | Escherichia coli E110019 | 340186 | 681 | 0.00067 |
| 12-benchmark-Complex | Escherichia coli O10:H32 | 2603836 | 497 | 0.00049 |
| 12-benchmark-Complex | Escherichia coli O18:H1 | 2126982 | 5670 | 0.0056 |
| 12-benchmark-Complex | Escherichia coli O15:H11 | 2048777 | 595 | 0.00059 |
| 12-benchmark-Complex | Escherichia coli O2:H6 | 1502658 | 3243 | 0.0032 |
| 12-benchmark-Complex | Escherichia coli M8 | 1392854 | 380 | 0.00038 |
| 12-benchmark-Complex | Escherichia coli VR50 | 941323 | 895 | 0.00088 |
| 12-benchmark-Complex | Escherichia coli O80:H26 | 2605620 | 1783 | 0.00176 |

| | | | | |
|---|---|---|---|---|
| 12-benchmark-Complex | Escherichia coli O139:H28 | 1603259 | 611 | 0.0006 |
| 12-benchmark-Complex | Escherichia coli PCN061 | 1358422 | 1019 | 0.00101 |
| 12-benchmark-Complex | Escherichia coli O6:H16 | 1446746 | 1498 | 0.00148 |
| 12-benchmark-Complex | Klebsiella pneumoniae subsp. pneumoniae | 72407 | 1379 | 0.00136 |
| 12-benchmark-Complex | Shigella dysenteriae 1617 | 754093 | 375 | 0.00037 |
| 12-benchmark-Complex | Shigella flexneri 2a | 42897 | 4492 | 0.00443 |
| 12-benchmark-Complex | Citrobacter rodentium ICC168 | 637910 | 295 | 0.00029 |
| 12-benchmark-Complex | Citrobacter amalonaticus Y19 | 1261127 | 79 | 0.00008 |
| 12-benchmark-Complex | Salmonella enterica subsp. enterica | 59201 | 6199 | 0.00612 |
| 12-benchmark-Complex | Salmonella enterica subsp. salamae | 59202 | 11 | 0.00001 |
| 12-benchmark-Complex | Xanthomonas euvesicatoria pv. alfalfae | 359387 | 181 | 0.00018 |
| 12-benchmark-Complex | [Ruminococcus] gnavus ATCC 29149 | 411470 | 1298 | 0.00128 |
| 12-benchmark-Complex | [Clostridium] scindens ATCC 35704 | 411468 | 44 | 0.00004 |
| 12-benchmark-Complex | [Clostridium] hylemonae DSM 15053 | 553973 | 12 | 0.00001 |
| 12-benchmark-Complex | [Clostridium] saccharolyticum WM1 | 610130 | 34 | 0.00003 |
| 12-benchmark-Complex | [Clostridium] sphenoides JCM 1415 | 1297793 | 25 | 0.00003 |
| 12-benchmark-Complex | Roseburia intestinalis L1-82 | 536231 | 61 | 0.00006 |
| 12-benchmark-Complex | Hungatella hathewayi WAL-18680 | 742737 | 58 | 0.00006 |
| 12-benchmark-Complex | Desulfitobacterium dichloroeliminans LMG P-21439 | 871963 | 11 | 0.00001 |
| 12-benchmark-Complex | Veillonella parvula HSIVP1 | 1316254 | 896 | 0.00088 |
| 12-benchmark-Complex | Rothia mucilaginosa DY-18 | 680646 | 250 | 0.00025 |
| 12-benchmark-Complex | Microbacterium testaceum StLB037 | 979556 | 12 | 0.00001 |
| 12-benchmark-Complex | Streptomyces lividans 1326 | 1200984 | 116 | 0.00011 |
| 12-benchmark-Complex | Eggerthella lenta DSM 2243 | 479437 | 59 | 0.00006 |
| 12-benchmark-Complex | Treponema pedis str. T A4 | 1291379 | 17 | 0.00002 |
| 12-benchmark-Complex | Human adenovirus 5 | 28285 | 270327 | 0.26674 |
| 12-benchmark-Complex | Human adenovirus 1 | 10533 | 31478 | 0.03106 |
| 12-benchmark-Complex | Severe acute respiratory syndrome coronavirus 2 | 2697049 | 41 | 0.00004 |
| 12-benchmark-Complex | Staphylococcus phage Andhra | 1958907 | 17 | 0.00002 |
| D1-downsampled-F41-2M | Xanthomonas euvesicatoria pv. alfalfae | 359387 | 11122 | 0.11044 |
| D1-downsampled-F41-2M | Streptomyces lividans 1326 | 1200984 | 20779 | 0.20634 |
| D1-downsampled-F41-2M | Mycobacterium tuberculosis variant caprae | 115862 | 27 | 0.00027 |
| D1-downsampled-F41-2M | Simian adenovirus 18 | 909210 | 16581 | 0.16464 |
| D1-downsampled-F41-2M | Simian adenovirus 16 | 1715778 | 22067 | 0.21912 |
| D1-downsampled-F41-2M | Simian adenovirus 1 | 310540 | 18913 | 0.18781 |

| | | | | |
|---|---|---|---|---|
| D1-downsampled-F41-2M | Human adenovirus 5 | 28285 | 202 | 0.00201 |
| D1-downsampled-F41-2M | Simian mastadenovirus WIV19 | 1923976 | 2654 | 0.02636 |
| D1-downsampled-F41-2M | Simian adenovirus 25 | 175567 | 1212 | 0.01204 |
| D1-downsampled-F41-2M | Human adenovirus B2 | 565303 | 1607 | 0.01596 |
| D1-downsampled-F41-2M | Simian adenovirus DM-2014 | 1560346 | 1356 | 0.01346 |
| D1-downsampled-F41-2M | Bat mastadenovirus WIV9 | 1788436 | 879 | 0.00873 |
| D1-downsampled-F41-2M | Human adenovirus 54 | 651580 | 858 | 0.00852 |
| D1-downsampled-F41-2M | Tree shrew adenovirus 1 | 47680 | 830 | 0.00824 |
| D1-downsampled-F41-2M | Simian adenovirus 49 | 995022 | 1332 | 0.01323 |
| D1-downsampled-F41-2M | Simian adenovirus 20 | 585059 | 133 | 0.00132 |
| D1-downsampled-F41-2M | Fowl aviadenovirus 5 | 172861 | 59 | 0.00059 |
| D1-downsampled-F41-2M | Staphylococcus phage Andhra | 1958907 | 93 | 0.00092 |
| D2-downsampled-F41-200K | Xanthomonas euvesicatoria pv. alfalfae | 359387 | 2004 | 0.11143 |
| D2-downsampled-F41-200K | Streptomyces lividans 1326 | 1200984 | 3814 | 0.21199 |
| D2-downsampled-F41-200K | Simian adenovirus 18 | 909210 | 3031 | 0.16846 |
| D2-downsampled-F41-200K | Simian adenovirus 16 | 1715778 | 3947 | 0.21942 |
| D2-downsampled-F41-200K | Simian adenovirus 1 | 310540 | 3438 | 0.1911 |
| D2-downsampled-F41-200K | Human adenovirus 5 | 28285 | 27 | 0.00151 |
| D2-downsampled-F41-200K | Simian mastadenovirus WIV19 | 1923976 | 483 | 0.02689 |
| D2-downsampled-F41-200K | Simian adenovirus 25 | 175567 | 230 | 0.01279 |
| D2-downsampled-F41-200K | Simian adenovirus DM-2014 | 1560346 | 258 | 0.01439 |
| D2-downsampled-F41-200K | Human adenovirus 54 | 651580 | 164 | 0.00912 |
| D2-downsampled-F41-200K | Bat mastadenovirus WIV9 | 1788436 | 163 | 0.00906 |
| D2-downsampled-F41-200K | Tree shrew adenovirus 1 | 47680 | 134 | 0.00745 |
| D2-downsampled-F41-200K | Simian adenovirus 49 | 995022 | 247 | 0.01376 |

| | | | | |
|---|---|---|---|---|
| D2-downsampled-F41-200K | Simian adenovirus 20 | 585059 | 16 | 0.00089 |
| D2-downsampled-F41-200K | Fowl aviadenovirus 5 | 172861 | 12 | 0.00067 |
| D2-downsampled-F41-200K | Staphylococcus phage Andhra | 1958907 | 19 | 0.00106 |
| D3-downsampled-F41-20K | Xanthomonas euvesicatoria pv. alfalfae | 359387 | 196 | 0.11708 |
| D3-downsampled-F41-20K | Streptomyces lividans 1326 | 1200984 | 376 | 0.22419 |
| D3-downsampled-F41-20K | Simian adenovirus 18 | 909210 | 309 | 0.18404 |
| D3-downsampled-F41-20K | Simian adenovirus 16 | 1715778 | 422 | 0.2518 |
| D3-downsampled-F41-20K | Simian adenovirus 1 | 310540 | 185 | 0.11024 |
| D3-downsampled-F41-20K | Simian adenovirus 25 | 175567 | 30 | 0.01821 |
| D3-downsampled-F41-20K | Simian mastadenovirus WIV19 | 1923976 | 59 | 0.03514 |
| D3-downsampled-F41-20K | Simian adenovirus DM-2014 | 1560346 | 26 | 0.01549 |
| D3-downsampled-F41-20K | Human adenovirus 54 | 651580 | 19 | 0.01132 |
| D3-downsampled-F41-20K | Simian adenovirus 49 | 995022 | 41 | 0.02476 |
| D3-downsampled-F41-20K | Bat mastadenovirus WIV9 | 1788436 | 13 | 0.00774 |
| D4-downsampled-F41-2K | Xanthomonas euvesicatoria pv. alfalfae | 359387 | 21 | 0.17146 |
| D4-downsampled-F41-2K | Streptomyces lividans 1326 | 1200984 | 39 | 0.32047 |
| D4-downsampled-F41-2K | Simian adenovirus 18 | 909210 | 24 | 0.19355 |
| D4-downsampled-F41-2K | Simian adenovirus 16 | 1715778 | 39 | 0.31452 |
